# Analysis of 3D epithelial tissue packing reveals correlated patterns of cell height and skewing

**DOI:** 10.64898/2026.01.14.699560

**Authors:** Guillaume Pernollet, Quentin Vagne, Juan Manuel García-Arcos, Claire A. Dessalles, Guillaume Salbreux, Aurélien Roux

**Affiliations:** Department of Biochemistry, University of Geneva, CH-1211 Geneva, Switzerland; Department of Genetics and Evolution, University of Geneva, CH-1211 Geneva, Switzerland; currently at Swiss Institute for Experimental Cancer Research (ISREC), School of Life Sciences, Swiss Federal Institute of Technology Lausanne (EPFL), Lausanne, Switzerland; currently at Université Claude Bernard Lyon 1, CNRS, Institut Lumière Matière, UMR5306, F-69100, Villeurbanne, France

**Keywords:** Cell volume, height fluctuation, epithelial monolayer, spherical confinement, mechanical environment, skewing, scutoids

## Abstract

Epithelia are tissues which envelop organs, exchange with and protect from their environment. The tight packing of cells within an epithelium guarantees its cohesiveness and impermeability, essential to its functions. Yet, how cells spatially arrange in three dimensions within an epithelium, and how the three-dimensional cell packing responds to geometrical constraints, is not fully understood. In a proliferating epithelium, cellular volumes vary from cell to cell, notably due to cell growth. At the same time, epithelia tend to display a smooth, continuous apical surface, indicating that cell shape determinants are spatially coupled. It is unclear how these and other factors modulate cell shape variability within the epithelium. Here, we segmented three-dimensional cell shapes of MDCK epithelia, grown in hollow spheres of alginate. With this assay, we could modulate the substrate adhesive properties, tissue curvature, and cell area density. We observed that as the tissue proliferates and cell density increases, average cell volume decreases. In contrast, cell height is relatively conserved over time, with an average value sensitive to the rigidity, curvature and adhesive strength of the substrate. We identified large and spatially correlated fluctuations in cell skewing, defined as the relative difference between apical and basal area that do not arise from curvature. Skewing is associated to spatial patterns of tilted cells whose apico-basal axis deviates from orthogonality to the substrate. Surprisingly, cell skewing and cell height are correlated, indicating internal rules for three-dimensional cell shapes. Altogether, our study identifies unexpected patterns of three-dimensional cell shape variation within a proliferative epithelium.

## Introduction

Tissues’ functions are linked to their shape and mechanical properties (1–3). Epithelial cells adopt roughly polyhedral shapes, typically tightly packed, forming cohesive layers to ensure tissue’s barrier function (4). Disruption in this cell organization leads to severe loss of function (5–8).

To understand epithelial geometry and mechanics, theoretical tools like vertex models (9–11) have been developed, which describe cells as polyhedra and introduce energy terms associated with cell surfaces and perimeters. In two dimensions, vertex models have been shown to account for tessellation properties of the apical side of epithelia with a limited number of parameters (9, 12). In three dimensions, the problem becomes substantially more complex, and three-dimensional vertex models often assume homogeneous volume and vertical lateral membranes (9), neglecting the possibility of tilt or more complex deformations along the apico-basal axis.

Cell volume and cell aspect ratio (AR, defined as cell width divided by height) are two simple descriptors of three-dimensional cell shape. Following perturbations on time scales of minutes, cells tend to conserve their volume. In this case, local changes in epithelium area, as invoked in tissue stretching (13) or tissue compression (14), are compensated for by cells changing their aspect ratio to preserve volume. Alternatively, during development, cells have been reported to lengthen along the apical–basal axis, while preserving their volume (15). Epithelial cells grown on a substrate with sinusoidal corrugations have been shown to modulate their aspect ratio to smoothen the apical surface, giving rise to spatial patterns of aspect ratio (16). Finally, on time scales much longer than the cell division time, average cell volume (17) and cell area (18) have been reported to decrease with increasing cell density.

However, cell volume and aspect ratio are still limited descriptors of three-dimensional cell shape, notably because they ignore differences in organization between apical and basal sides of epithelia. Such differences have been reported and attributed either to changes in cell area along the apical-basal axis (19), or to the magnitude of curvature. Gradients of curvature (20) and curvature anisotropy (21) can induce neighbor exchange from apical to basal side, analogous to a T1 transition but along the apical-basal axis, leading to anisotropic cell shapes sometimes referred to as scutoids.

In this study, we aimed at evaluating the impact of different constraints on three-dimensional cell shapes within an epithelial packing. To control curvature, mechanical properties of the substrate and adhesion, we encapsulated epithelial cells in hollow spheres of alginate. Adhesion to the inner or outer surface of the hollow sphere was modulated by addition of Matrigel or RGD-peptide to the alginate. We then developed an analysis pipeline to segment and analyze cell shape in three dimensions.

With this approach, we show that as cells proliferate and the average cell density increases, the average cell volume decreases. The average cell height was conserved over time, with a value that depended strongly on the rigidity and adhesive strength of the substrate. Looking at spatial patterns of cell height and volume, we found that the volume fluctuations are largely uncorrelated and volumes vary strongly between neighbors, while cell height is spatially correlated, with a persistence length of a few neighbors. We further observed that height fluctuations at long distances robustly correlate with cell skewing and tilt. Such 3D changes in cell organization additionally correlate with the observation of highly conical cells and scutoids, even on flat substrate, therefore unrelated to curvature.

## Results

### 1) Cell encapsulation as a tool to study tissue organization

To investigate the impact of geometrical constraints on tissue organization, we used Madin-Darby canine kidney II (MDCK II) cells to grow epithelial monolayers on the inner wall of hollow alginate microspheres, with diameters ranging between 80 and 200 µm. (hereafter referred to as capsules). These capsules were produced by the co-extrusion of three fluid flows using a microfluidic device previously developed in our laboratory. (Figure 1A) (22).

**Figure 1:**
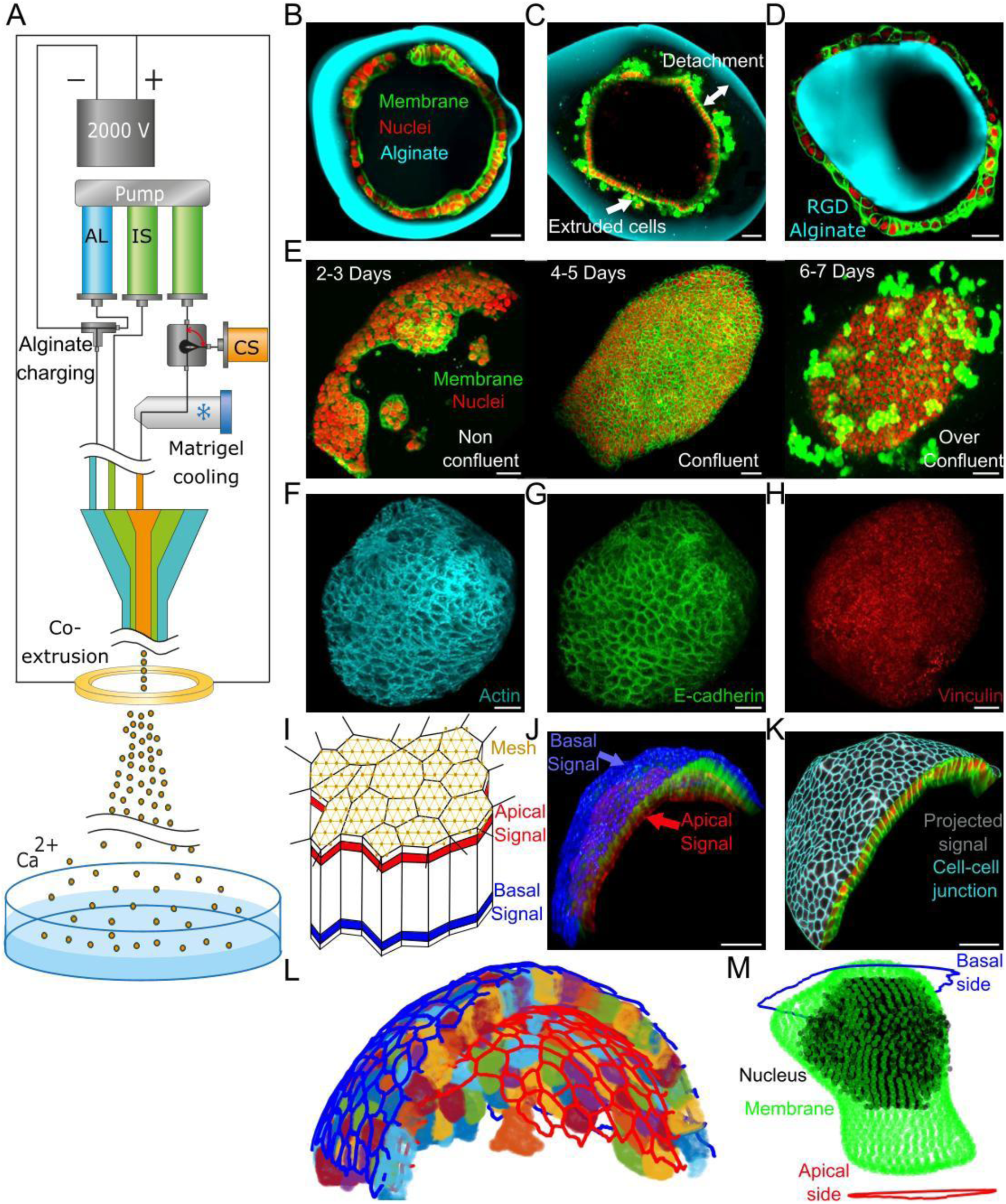
Development of a segmentation pipeline for 2D+ tissue tessellation study. Schematic of the experimental coextrusion capsule-making setup (A). Tissue/capsule interactions showing confinement, contractile detachment, and convex curvature substrate (cells grown on the outer surface), scale bar 30um (B-D). Timeline of growth of a WT MDCK monolayer within capsules, scale bar 30um (E). Max intensity projection of Actin, cadherin and vinculin staining for WT MDCK, scale bar 30um (F-H). Schematic of MorphographX segmentation: Signal used for basal segmentation (blue) and apical segmentation (red) is projected onto a surface mesh (I). Membrane signal used for basal segmentation (blue) and apical segmentation (red), scale bar 30um (J). Example of 2D+ segmentation of the basal side, scale bar 30um (K). Example of 2D+ segmentation of apical side (red), basal side (blue) and the 3D cell segmentations performed using Limeseg (L). Example of a single cell segmentation, combining MorphographX (apical/Basal segmentation), starDist (nuclei segmentation) and Limeseg (volume segmentation) (M).

This method allows to study tissue growth under 3D elastic confinement and without boundary effects because of the spherical geometry (22–24)(Figure 1B). Nutrients and growth factors are accessible to the encapsulated cells thanks to the use of permeable alginate gels (25). In this environment, cells grow over multiple days, until tissues detached from the inner surface, a process accompanied by massive cell extrusion, forming a smaller structure within the capsule (Figure 1C). The capsule diameters - and therefore tissue curvatures - can vary from 50 µm to 200 µm, by modulating the microfluidic device’s nozzle diameter and the exit flow rates. As cells do not adhere on alginate, we routinely use a layer of Matrigel to promote cells adhesion to the inner surface of capsules(22, 25). As Matrigel’s adhesive properties can be hardly changed, cell adhesion can be modulated using alginate covalently labelled with the RGD peptide, a peptide binding to integrins, and changing the fraction of RGD-alginate in the alginate mix. The use of RGD-alginate also meant that the entire capsule was adhesive, allowing us to grow tissues on the outer surface (Figure 1D).

As capsules allowed us to pinpoint the moment tissues reached confluency, covering the entire capsule surface, we decided to sort tissues into three broad categories: i-non-confluent tissues, representing the majority of tissues between the encapsulation point and the 3rd day (Figure 1E left), ii-confluent tissues representing the majority of tissues between the 4th and 5th day (Figure 1E center), and iii-over-confluent tissues past the 5^th^ day mark (Figure 1E right), defined by the apparition of detachment and cell extrusion.

To extract the position of cells and identify them, we labelled nuclei with Hoechst, and to extract the cell shapes, we used Cell-Mask to stain the plasma membrane. Changes of acto-myosin organization were evaluated by staining cells with phalloidin tagged with Alexa Fluor and changes in cell adhesion were observed through immuno-staining for E-cadherin and Vinculin (Figure 1F-H).

Lastly, in addition to the wild-type MDCK II cells (WT MDCK), the membranes of which can be labelled by chemicals, we used an MDCK II cell line stably expressing Myr-Palm GFP, where a GFP is fused to a myristoylation and palmitoylation domain to anchor it to the plasma membrane, and H2B-mCherry, a histone marker that labels the nuclei. We refer to this cell line as MP MDCK in the following. Initially developed to allow lifetime imaging to track the cell membrane and nuclei, this cell line displays different organization and contractile properties that WT MDCK, compatible with MP MDCK cells being more contractile (25). Nonetheless, these cells still grew and covered the capsule surface, before detaching after multiple days (Supp Fig 1A).

### 2) Development of an analysis pipeline for tissue organization

To extract cell geometrical characteristics, we developed a unique analysis framework, where cell nuclei and surfaces are first segmented in 3D using a custom pipeline (Figure 1 I-K, Materials and Methods). Extracting geometrical features of the apical and basal faces from 3D segmentation of epithelial cells is difficult and unreliable. We therefore segmented apical and basal surfaces of the epithelium independently, and then apical and basal segments were assigned to specific cells in the tissue using positional information of the nucleus. These three segmentations were processed through a custom Matlab code to get a full 3D segmentation of the entire capsule (Figure 1L) and extract over twenty 2D/3D geometrical parameters (Table 1), including the curvature computed from the basal mesh. These parameters were obtained for each cell (Figure 1M) and their neighbors, allowing us to generate auto- and cross-correlation maps (Supp Fig 1B-C).

**Table 1:**
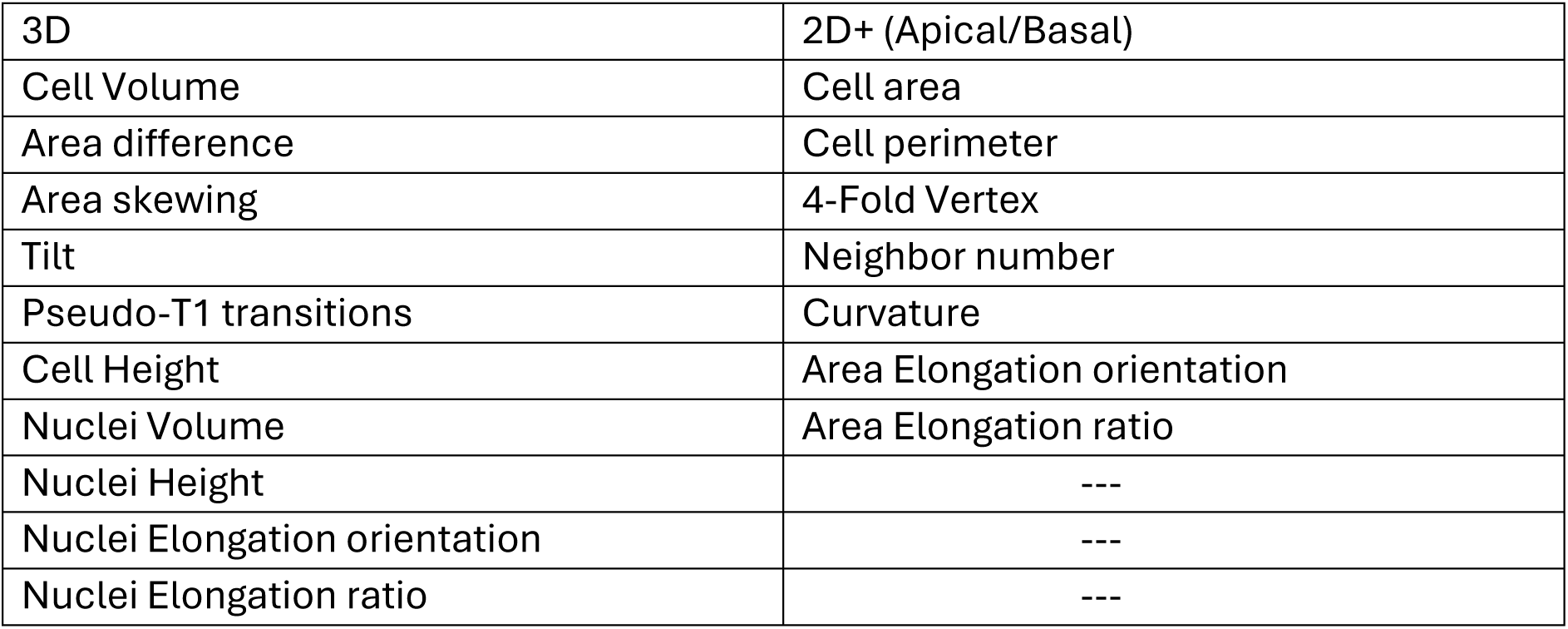

This tool, in combination with the cell encapsulation technique, allowed us to carry an in-depth investigation of tissue tessellation with a 3D perspective on the epithelium organization, and to follow how this organization evolves as the cell density increases.

### 3) Cells undergo volume changes and remodeling with increasing density

We first decided to investigate how epithelial cell volume changed with varying cell densities. For this, we imaged and segmented different encapsulated tissues grown from 2 to 7 days post-encapsulation (Figure 2A). We evaluated cell volume and cell density, obtained from the inverse of the cell basal area. Cell volume continuously decreased with time, until most cells reached a volume round 300 µm^3^ (Figure 2B-C). We found an overall robust anticorrelation between cell volume and cell density, with the volume undergoing a six-fold decrease, approaching a minimum value with increasing density (Figure 2D), while always following a log-normal distribution (Figure 2E). These findings support that epithelial cells adapt their volume to cope with increasing density, consistent with previous work (17). We then wondered whether the volume of nuclei followed the same trend. Average nuclear and cell volumes were linearly related to each other with a slope less than one, indicating that nuclei occupy a larger relative fraction of cell volume when the cell volume decreases (Figure 2F-G), as multiple recent studies reported (17, 28).

**Figure 2.**
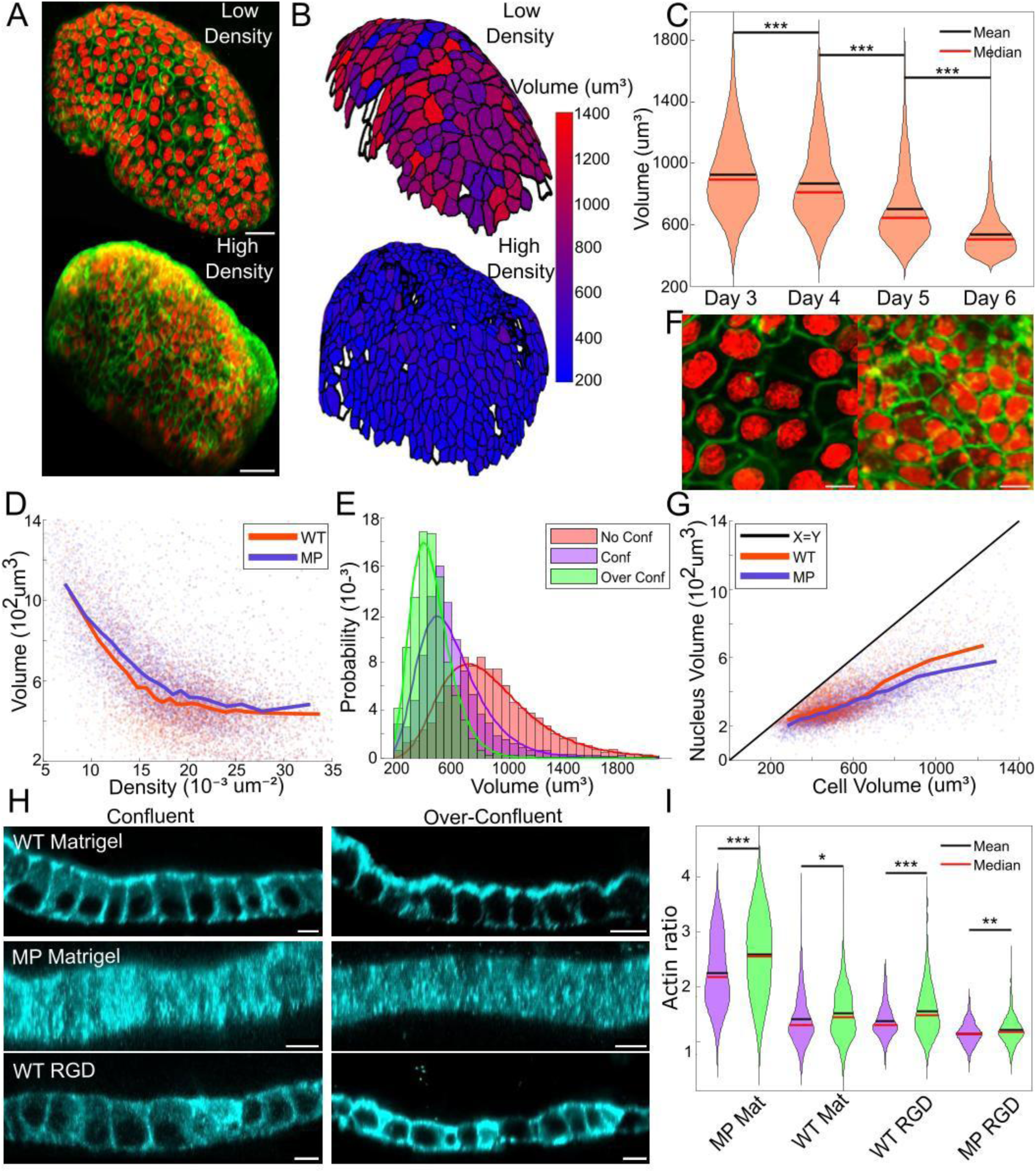
Volume behavior upon geometrical confinement. WT MDCK tissue stained for membrane (green) and nuclei (red) at 2 days post-encapsulation (top) and 6 days post-encapsulation (bottom), scale bar 30um (A). Cell volume at 2 days post-encapsulation (top) and 6 days post-encapsulation (bottom) (B). Cell volume as a function of days post-encapsulation, Matrigel grown WT, (N=10 tissue per condition) (C). Single cell volume as a function of density for Matrigel-grown WT (N=20) and MP (N=27) tissues (D). Distribution of volume for Matrigel grown WT, fitted with lognormal distributions, for the non-confluent case (N=13), confluent case (N=11) and overconfluent case (N=12) (E). WT tissue stained for membrane (green) and nuclei (red) at 2 days post-encapsulation (left) and 6 days post-encapsulation (right), scale bar 10um (F). Nuclei volume for WT and MP as a function of cell volume (WT: N=20, MP: N=27) (G). Actin signal for Matrigel grown WT (top), Matrigel grown MP (center) and RGD grown WT (bottom) at confluency (left), and over-confluency (right), scale bar 10um (H). Actin ratio between lateral (purple) and apical (green) sides for WT and MP Matrigel and RGD grown tissue (N=3 tissues for each condition) (I). Thick lines: average of bins containing equal numbers of elements (D, G). *: p<0.05, **: p< 0.01, ***: p<0.001.

We wondered if the decrease in volume was associated with reorganization of the actin cytoskeleton and adhesion sites, as both are essential actors of cell packing. For WT MDCK grown on Matrigel, actin organization was highly polarized with a bright apical actin staining (Figure 2H), and low basolateral intensities. WT MDCK cells grown on RGD displayed much less differences between apical and basolateral intensities (Figure 2H-I). On RGD-alginate, stress fibers were present at the basal side, and absent on Matrigel (Supp Fig 2A). This may reflect the higher rigidity of RGD-Alginate over Matrigel, as stress fibers are known to appear on rigid substrates (29). For MP MDCK cells on Matrigel, actin appeared much less polarized than for WT MDCK. Furthermore, actin appeared to accumulate more at the lateral junctions on RGD grown tissues (Figure 2H-I) and never displayed stress fibers regardless of the substrate (Supp Fig 2B). These observations may account for the fact that MP MDCK cells have a more homogeneous, less polarized contractility, whereas WT MDCK can hyperpolarize their contractility to the apical face but also reorganize contractility depending on the basal substrate. We also observed that the polarization of actin, which means the intensity difference of actin staining on the apical side to basolateral sides increased with increasing cell density in all tissues (Figure 2I).

In confluent tissues, while cadherin staining didn’t display strong differences between different cell types and substrates (Supp Fig 2C-D), it changed with cell density. For Matrigel grown tissues, cadherin delocalized from the cell-cell junctions and accumulated in the cytoplasm as the tissue became overconfluent, while vinculin displayed a stronger basal localization at high cell density (Supp Fig 2E), particularly in WT MDCK.

Altogether, our results show that the volume decreases with time and increasing tissue density, while cell cortex organization is remodeled. This led us to wonder how other geometrical quantities such as cell height and area evolved and correlated.

### 4) Substrate properties control the average cell height independently of cell volume

We next studied the correlations between average cell height and average cell volume within capsules at different time points (Figure 3A). We observed that the average height was roughly independent of cell volume, in spite of the previously mentioned large volume variations (Figure 3B). Despite this global independence to cell volume, cell height could still vary across tissues or within a single tissue. Indeed, tissues grown on RGD alginate presented much smaller height than Matrigel grown tissues (Figure 3C-D). We hence wondered which properties of the substrate could impact the height. As curvature (16), substrate rigidity (30) and substrate adhesive properties (31) have already been observed to impact tissue organization, we decided to further investigate their impact on tissue height.

**Figure 3:**
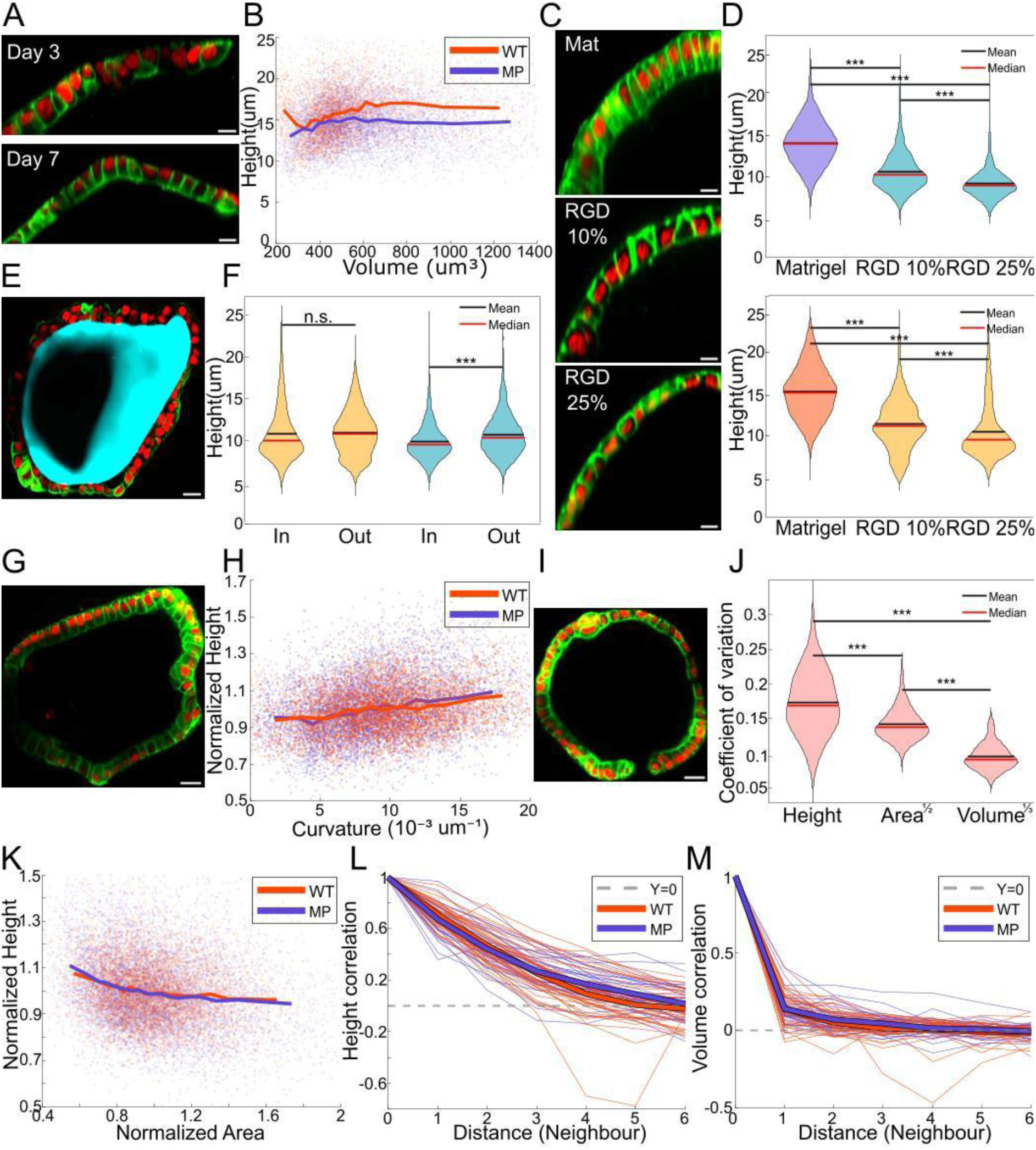
Height control by mechanical parameters Lateral slice of Matrigel grown WT tissue stained for membrane (green) and nuclei (red) at 3 days post-encapsulation (top) and 7 days post-encapsulation (bottom), scale bar 10um (A). Individual cell height as a function of cell volume for WT MDCK and MP MDCK (WT: N=20 capsules, MP: N=27 capsules) (B). Lateral slice of RGD grown WT and Matrigel grown WT tissue, at confluence, stained for membrane (green) and nuclei (red), scale bar 10um (C). Impact of adhesion molecules on tissue height for MP tissues (top) (Matrigel N=27, RGD 10%: N=10, RGD 25%: N=13) and for WT tissues (bottom) (Matrigel: N= 20, RGD 10%: N=5, RGD 25%: N=12) (D). RGD grown WT tissue, under convex curvature, stained for membrane (green) and nuclei (red), scale bar 20um (E). Quantification of cell height for RGD grown WT and MP tissues, under concave (in) and convex (out) curvature (WT, RGD 25% In N= 13, RGD 25% Out N= 9, MP RGD 25% In N= 12, RGD 25% Out N= 4) (F). WT Matrigel grown tissue experiencing different curvatures imposed by capsule shape, scale bar 20um (G). Impact of curvature on tissue height for Matrigel grown tissues (WT: N=20, MP: N=27) (H). MP MDCK tissue displaying large differences in height within a single tissue, stained for membrane (green) and nuclei (red), scale bar 20um (I). Coefficient of variation for Height, Area^1/2^ and Volume^1/3^ for Matrigel grown WT (N=20) (J). Single cell normalized Height as a function of normalized area, for Matrigel grown WT and MP tissue (WT: N=20, MP: N=27) (K). Height autocorrelation as a function of the distance, quantified in cell number, for Matrigel grown WT and MP tissue (WT: N=20, MP: N=27) (L).Volume autocorrelation as a function of the distance, quantified in cell number, for Matrigel grown WT and MP tissue (WT: N=20, MP: N=27) (M).Thick lines: average of bins containing equal numbers of elements (B,H,K), or average at each cellular distance (L, M). *: p<0.05, **: p< 0.01, ***: p<0.001.

As stronger adhesion was shown to lead to more cell spreading and hence smaller height (32), we first tested for the effect of substrate adhesive properties, by changing the fraction of Alginate-RGD. Increasing Alginate-RGD from 10% to 25% (v/v) in the alginate mix led to a decrease in height, which was more pronounced for WT MDCK than for MP MDCK (Figure 3D). MP MDCK may be less sensitive to changes in substrate adhesive properties because these cells do not form basal stress fibers, possibly reducing their ability to spread.

MDCK cell height has been reported to change at substrate rigidities around 1kPa (30). To test the effect of substrate rigidity, MDCK epithelia were grown on flat commercial hydrogels with rigidities of 0.5kPa and 4kPa subsequently coated with fibronectin. Surprisingly, we observed that WT tissues had higher values of cell height on softer tissues, while it was the opposite for MP tissues (Supp Fig 3A). As with encapsulated tissues, we observed no correlation between cell height and volume at 4kPa but observed a non-negligible correlation at 0.5kPa for WT tissues (Supp Fig 3B-D).

To rationalize these observations, we turn to a simple theoretical argument. We assume that cell shape arises from the minimization of an effective mechanical energy *E*. We consider the following contributions: a uniform cortical surface tension acts on all cellular (apical, basal, lateral) interfaces (*σ*_*c*_), an additional surface tension acts on the apical surface (*σ*_*a*_) and an adhesion energy density (*χ* > 0) acts as a negative surface tension on the basal side (Supp Fig 3E). Writing the corresponding expression for *E*, imposing the cell volume and minimizing for cell height leads to:

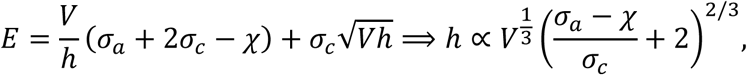

with *V* the cell volume and *h* the cell height. If we now assume that cells respond to a softer substrate by lowering their overall cortical contractility *σ*_*c*_, leaving adhesion energy and apical contractility unchanged, then the sign of *σ*_*a*_ − *χ* controls how height responds to substrate rigidity. For WT tissues, if apical contractility dominates over adhesion (*σ*_*a*_ > *χ*), cell height increases on softer substrates. MP MDCK cells’ actin cortex appears less polarized (Figure 2H): in this case, adhesion may dominate over apical contractility (*σ*_*a*_ < *χ*), leading to a decrease in cell height on softer substrates.

To further test this picture, we perturbed contractility by treating encapsulated tissues with the non-muscle myosin II inhibitor blebbistatin. Epithelia were grown until confluency and imaged using actin staining (Supp Fig 3F) before adding 10µM of blebbistatin (25). One hour after blebbistatin treatment, tissue height of WT MDKC cells was not affected, whereas the height of MP tissues decreased (Supp Fig 3G). This can be rationalized with the model introduced above: assuming blebbistatin treatment affects all tension terms equally, in WT cells, the ratio of apical to cortical contractility (*σ*_*a*_/*σ*_*c*_) dominates, and height changes should be small. For MP cells, however, the adhesion to cortical contractility ratio (*χ*/*σ*_*c*_) dominates and the height is expected to decrease.

We then took advantage of the irregular shapes of capsules to ask whether substrate curvature can impact tissue height. We used RGD alginate capsules, as cells can adhere onto the inner surface, resulting in positive (concave) curvature, or onto the outer surface, resulting in negative (convex) curvature (Figure 3E; we define curvature as positive if the apical surface is more highly curved than the basal surface). We reasoned that due to apical contractility, cell height might increase for cells adhering to the inner surface, and decrease in cells adhering to the outer surface, as the apical tension would couple to curvature to result in a force pulling inward (16). However, we did not observe a significant drop in height in cells grown on the outer surface, nor an increase in cells grown on the inner surface (Figure 3F).

To investigate further the effect of curvature on tissue height, we asked how cell height depends on the tissue local curvature, limiting ourselves to positive curvatures (Figure 3G, cell height is normalized with respect to cell height average within each tissue). A robust and significant correlation between cell height and curvature was observed for both WT and MP Matrigel grown tissues (Figure 3H). It was also present in RGD grown tissues (Supp Fig 3H). Thus, tissue curvature and cell height have a complex, non-monotonous relationship, as both increasing and decreasing curvature can lead to increased cell height.

### 5) Cell height is more variable than cell size, but correlated in space

Although we found that substrate properties influence average tissue height, cell height nevertheless presented large variations between tissues in a given condition, and large variations between cells within a single tissue (Figure 3I). To quantify cell height variability relative to other cell dimensions, we also calculated the cell width *w*, defined as:

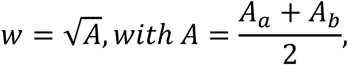

where *A*_*a*_ and *A*_*b*_are the apical and basal surface areas and *A* is the average between the two (simply called “area” then on). Despite the observed large density variations, we found that *h* and *w* showed similar coefficients of variations for Matrigel grown tissues (Supp Fig 3I). In the case of RGD-grown tissues, the height fluctuated even more than the width, particularly for WT cells. Interestingly, when quantifying height fluctuations within single tissues, we found that the coefficient of variation was only slightly lower than the global inter-tissue coefficient of variation (Supp Fig 3J). This prompted us to study in more detail height fluctuations within individual tissues.

In addition to cell height *h* and cell width *w*, we evaluated the cell size *R* = *V*^1/3^ for a given cell volume *V*. Comparing variations in *h*, *w*, and *R*, we found that surprisingly, within a single tissue, cell size shows the smallest variability (Figure 3J), suggesting that volume fluctuations alone cannot explain the observed height and width fluctuations. We also reasoned that if cell shapes were fluctuating at fixed aspect ratio, but varying cell volume, cell area and height would be positively correlated. In contrast to this picture, a small anticorrelation between cell area and height can be observed (Figure 3K, Supp Fig 3K-L), irrespective of the tissue stage (non-confluent, confluent, over-confluent) and the substrate.

We then asked if cell height, although varying significantly across an epithelium, was spatially correlated. Indeed, the Pearson correlation of cell height between nearest neighbors was 0.66 for MP and 0.7 for WT, then decreased over a length scale of a few cells (Figure 3L). In contrast, cell volume did not show long range correlation (Figure 3M) and had a Pearson correlation of 0.14 for both cell types between nearest neighbors.

Overall, we concluded that although cell volume varied substantially over time, the overall cell size is less variable than cell width or cell height across the epithelium, at a given time point. Although more variable than cell size, it is cell height, but not cell volume, that is spatially correlated on a scale of a few cells.

### 6) Height fluctuations are associated to lateral junction tilt and cell skewing

We then asked whether the apical and basal sides exhibited differences in packing (Fig 4A). Calculating a unitless shape order parameter defined as 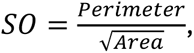 independently for apical (*SO_a_*) and basal (*SO_b_*) surfaces, we could estimate whether the apical and basal packings were close to a regular hexagonal arrangement (Supp Fig 4A). For a regular, crystalline hexagonal packing, *SO =* 3.722, with higher values indicating more disordered packings (33). *SO_a_* and *SO_b_* have similar statistical distributions (Supp Fig 4B) and were largely independent of single cell volume and area (*A*, averaged between apical and basal) across tissues (Supp Fig 4C-D). The distributions of area on the apical and basal sides were also similar (Supp Fig 5A).

**Figure 4:**
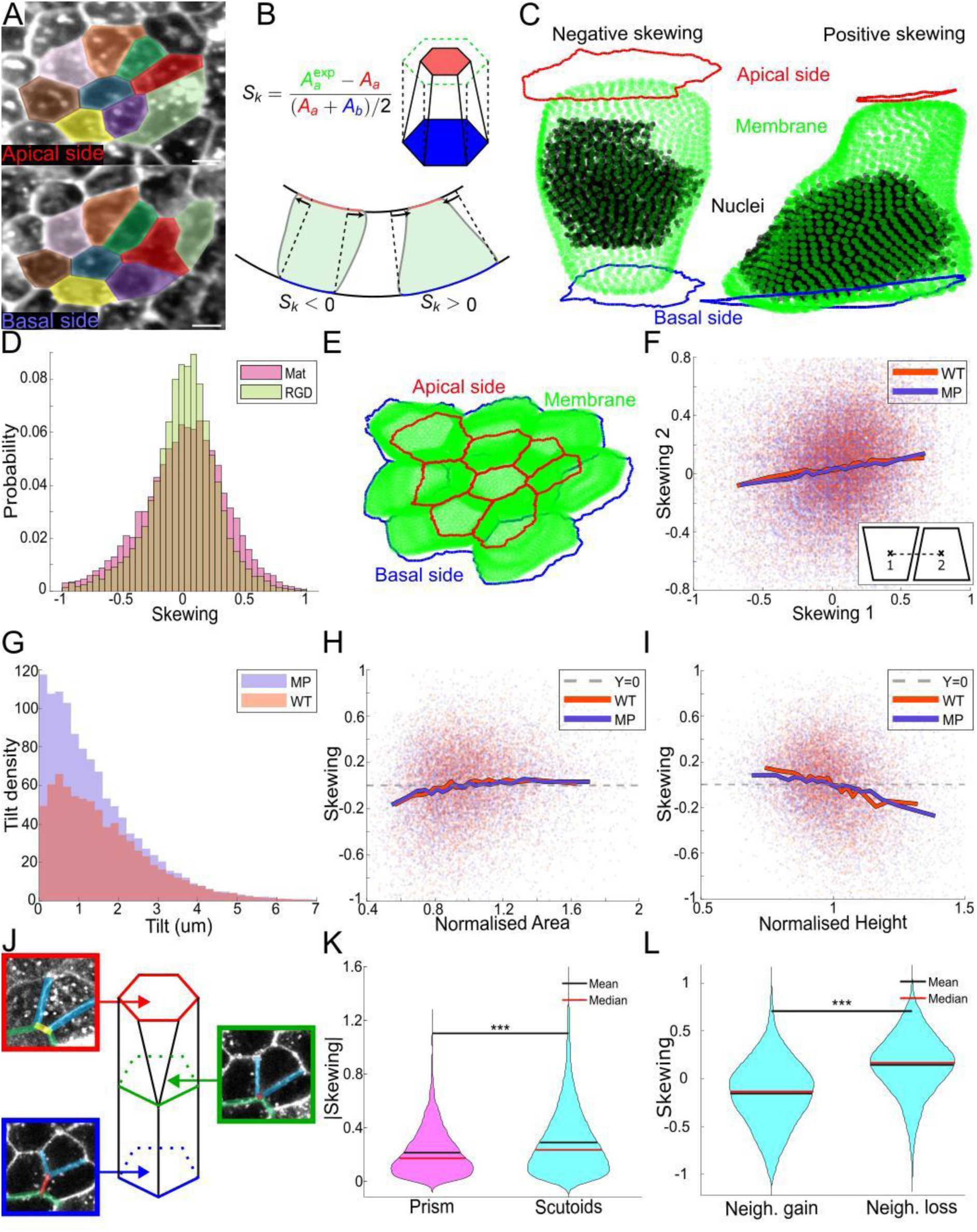
Apical/basal organization discrepancy. Apical (top) and basal (bottom) membrane organization for a WT MDCK tissue grown on Matrigel, scale bar 5um (A). Schematic illustrating how skewing is defined and computed (B). Example of cell displaying negative skewing (left) and positive skewing (right) (C). Distribution of cell skewing for Matrigel and RGD grown tissues (Matrigel: N=47, RGD: N= 41) (D). Example of local region of positively correlated cell skewing (E). Cell skewing vs skewing of the neighboring cell for Matrigel grown tissues (WT: N=20, MP: N=27) (F). Distribution of tilt for Matrigel grown MP and WT tissue (WT: N=20, MP: N=27) (G). Skewing vs normalized area ((*A*_*a*_ + *A*_*b*_)/2) (WT: N=20, MP: N=27) (H). Skewing vs normalized height (WT: N=20, MP: N=27) (I). Illustration of apical/basal T1 transition, showing a yellow junction on the apical side, progressively disappearing and replaced by a red junction on the basal side (J). Skewing norm distribution for prism-shaped cells and scutoid-shaped cells for Matrigel grown tissues (N=47) (K). Skewing of scutoids gaining a neighbor along their basal-to-apical axis, and of scutoids losing a neighbor along their basal-to-apical axis (N=47) (L). Thick lines: average of bins containing equal numbers of elements (F, H, I). *: p<0.05, **: p< 0.01, ***: p<0.001.

The average 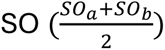 was dependent on the type of substrate (Supp Fig 4E-F), with tissues grown on stiff substrates (hard hydrogels and RGD-alginate) always being more disordered than tissues grown on soft substrates (soft hydrogels and Matrigel). Considering the model presented above, where cortical tension increases with rigidity, this hints that higher cortical tension leads to tighter packing. This goes in contrast with the usual assumption that apical tension in the main parameter controlling packing.

As *SO_a_*, *SO_b_*, apical and basal area distributions are similar at the population level, we asked about the relationship between apical area *A*_*a*_and basal area *A*_*b*_within the same cell. To address this point, we defined a new cell shape parameter called skewing, which we denote *Sk* (Figure 4B):

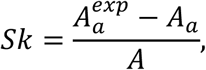

measuring the deviation of the measured apical area *A*_*a*_ from an expected area 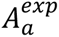 that depends on curvature and is equal to the basal area for a flat substrate. The expected area 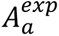 corresponds to the apical area predicted from the basal area, local mean curvature and tissue height, under the assumption that cell lateral junctions are normal to the substrate, and reads:

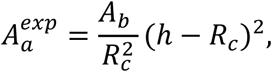

with *R*_*c*_the curvature radius, *h* the cell height and *A*_*b*_the measured basal area. Skewing is positive when the cell is more contracted apically than is expected from its basal area, and negative when the cell is less contracted (Figure 4C, Supp Fig.5B-C). *Sk* has lower and upper theoretical bounds -2 and 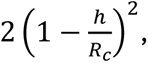 which correspond respectively to a vanishing basal and a vanishing apical area.

Experimental values of *Sk* are, however, largely confined between -0.5 and 0.5 (Figure 4D). We wondered about possible cell shapes giving rise to such magnitude of skewing. On a flat substrate, assuming an ideal conical shape and a skewing of 0.5, cell lateral junctions are tilted by an angle that depends on the cell aspect ratio. For columnar tissues 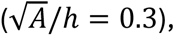 such a skewing magnitude can arise from lateral junctions tilted by just 2°, while for squamous tissues 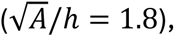 it would require a tilt of 15° (Supp Fig.5D). The 3D segmentations, however, reveal that cells are not ideal frustums: lateral junctions are not straight and appear more deformed near the nuclei (Figure 4C), as previously reported (19). At the global level, we observed in all epithelia at all densities that cells display a non-negligible degree of skewing (*Sk* ≠ 0, Figure 4D, Supp Fig 5E) even though the average of *Sk* vanishes. The skewing distribution appeared to be asymmetric, with more cells having positive skewing than negative skewing (Supp Fig 5F). The spread of the skewing distribution appeared independent from the cell type but was impacted by the substrate, the skewing distribution being slightly narrower on Matrigel than on RGD (Supp Fig 5G).

How are cells with different skewing packed together within an epithelium? A simple possible organization could involve an anti-correlation of skewing between nearest neighbors, where cells with roughly conical shapes would be compensated by neighboring inverted cones. To test this, we calculated the skewing correlation between neighboring cells and found, surprisingly, that the correlation was slightly positive (Figure 4E-F), indicating that cells with the same skewing orientation tend to group together. Such spatial skewing correlation should either induce local increase in curvature or cell tilt. Cell tilt occurs when the cell main axis, connecting the centers of mass of the apical and basal surfaces, are not aligned with the direction normal to the epithelial plane. To evaluate cell tilt, we calculated the vector connecting the barycenters of the apical and basal faces of cells. We then calculated the tilt as the distance between the intersection of the vector normal to the basal plane and the apical surface, and the apical barycenter. We observed that a high proportion of cells are indeed tilted (Figure 4G) and that the amplitude of tilt correlates with |*Sk*| (Supp Fig. 5H).

We then asked whether cell skewing could be related to other cellular properties. We found that nucleus position correlated with skewing, with positively skewed cells localizing their nuclei more basally (Supp Fig 5I) as expected from steric effects. Cell skewing was also weakly anti-correlated with cell volume (Supp Fig 5J) but independent from the shape parameter *SO* (Supp Fig 4G). Interestingly, WT and MP MDCK cells grown on flat hydrogel substrates had the same distribution of skewing than epithelia grown in capsules (Supp Fig 5K), indicating that cell skewing is not induced by curvature of the epithelium.

Surprisingly, we observed a correlation between cell skewing and cell area, with cells with larger areas exhibiting positive average skewing (Figure 4H); this correlation is observed at all cell densities (Supp Fig. 5L). Since cells with a small area have a larger height (Figure 3K), one would then expect an anti-correlation between cell skewing and cell height. Indeed, taller cells tend to be negatively skewed (Figure 4I). Thus, cells with larger height tend to have larger apical than basal area. We note that such a correlation is opposite to what would be intuitively expected from fluctuations in apical tension, which would simultaneously drive positive skewing (smaller apical areas) and increased cell height (to compensate for the volume decrease induced by this contraction). But consistently at the single tissue level, |*Sk*| increases roughly linearly with the average cell height (Supp Fig. 5M). We also noticed that WT cells are more skewed on soft substrates, where they are taller, while MP cells are more skewed on rigid substrates, where they are taller (Supp Fig 5K).

Finally, skewing showed a very small anti-correlation with substrate curvature (Supp Fig 5N). However, we observed that tissues with negative curvature had a broader distribution of skewing than those on positive curvature (Supp Fig 5O).

### 7) Skewing induces the formation of apico-basal T1 transition caused by area shrinkage

In three dimensions, epithelia have been reported to exhibit apico-basal T1 neighbor exchanges, a topological change sometimes referred to as a “scutoid” (Figure 4J). Such transitions have been previously associated with high curvature gradients (20, 34), curvature anisotropy (21), or pseudo-stratified epithelia (19), where they were attributed to the highly irregular shape of cells and distribution of nuclei position. This prompted us in investigating whether cell skewing could be associated with apico-basal T1 neighbor exchanges.

Interestingly, we observed such transitions in all epithelial tissues, including flat tissues. The fraction of cells showing at least one apico-basal T1 transition was similar across all conditions, around 50%, and was the same for flat and curved tissues (Supp Fig 6A). Consistent with this, cells participating in apico-basal T1 transitions were not on average in regions with higher curvature (Supp Fig 6B). These cells also had similar volume and area distributions than regular frustum cells (Supp Fig 6C-E). However, they were significantly more skewed, taller and more tilted (Figure 4K, Supp Fig 6F-G) supporting the idea that skewing is associated with apico-basal T1 transitions.

To investigate this in more detail, we looked at cell junctions. For a given cell exhibiting an apico-basal T1 transition, one or several junctions disappear from one side of the epithelium to the other (Figure 4J). We found that cells losing neighbors along the basal-to-apical axis had a positive skewing (smaller apical area), while cells gaining neighbors were negatively skewed (Figure 4L), supporting the hypothesis that scutoids emerged from skewing. We additionally observed that disappearing junctions tended to be shorter than traversing junctions (Supp Fig 6H).

Altogether, these results support the notion that local control of height combined with large differences of volume in neighboring cells causes skewing of epithelial cells. The local height control also explains why cells skewed with the same orientation appear to be grouped, forcing them to tilt. Also, skewing, by shrinking one face of the cells, induced the fusion of vertices and the disappearance of one neighbor from the smallest side, creating scutoidal cells.

## Discussion

In this study, we show that MDCK epithelial monolayers undergo drastic reduction in average cell volume and increase in cell density while proliferating inside alginate capsules (Figure 2A-D), independently from external confinement (Figure 2E). This is in contrast with the view that cell volume is preserved on average during tissue proliferation (35, 36), such that increasing cell density results in increasing cell height to preserve volume. Recently, other studies have hinted at cell area (18) and volume (17) decreasing as tissues reached confluency.

Although average cell height was largely independent from cell volume (Figure 3B), we found that cell height was sensitive to the substrate adhesive properties, its rigidity and its curvature (Figure 3D,3F). A simple model that combined uniform cortical tension, basal adhesion energy, and apical-specific tension was sufficient to explain the effects of varying substrate rigidity, substrate adhesive properties, and blebbistatin treatment; assuming that cells react to substrate rigidity by increasing their cortical tension while keeping their adhesion energy constant, as rigidity has been observed to impact cell polarity (37).

We identify skewing as a mode of three-dimensional cell deformation (32, 38, 39). Skewing is spatially correlated, shows similar distributions in tissues of different densities, and is negatively correlated to cell height. It would be interesting to explore whether fluctuations in interfacial cellular tensions, as well as volume fluctuations, can explain the origin of this correlation.

While our work highlights the complexity of the 3D organization of simple epithelial monolayers, it would be interesting to explore whether skewing, tilt or height differences influence processes such as cell division or contractile forces generated within the cell; such feedback loops between force-generating processes and cell geometry may be at the origin of the distribution of three-dimensional cell shapes within an epithelium.

## Acknowledgments

We thank Y. Ravichandran, I. De Meglio, M. Luciano, and S. Gabriel for helpful scientific discussions, as well as all members past and present of the Roux lab and Salbreux lab. We are grateful to I. De Meglio for training and assistance with alginate capsule preparation, and to M. Luciano and S. Gabriel. for discussions and training related to hydrogel production and mechanical characterization. A.R. acknowledges funding from the Swiss National Science Foundation (grant numbers #131003A_173087, #CRSII5_189996, and #310030_200793), the NCCR Chemical Biology, and the European Research Council Synergy grant #951324-R2-TENSION.

## Author contributions

G.P., Q.V., A.R. and G.S. designed the research. C.D. developed the protocol for RGD-alginate capsules. G.P. performed the experiments. G.P., Q.V. and G.S. conceived the data analysis protocol. G.P. performed the segmentations, developed the Matlab code for the analysis pipeline. G.P. and Q.V. analyzed the data. G.P., Q.V., G.S., J.M.G and A.R. discussed data interpretation and conceived the figures. G.P., Q.V., G.S. and A.R. wrote the paper, with inputs from all authors.

## Supplementary Information

## Methods

### Cell culture and stable cell line generation

Madin-Darby Canine Kidney II (MDCK-II) (ECACC, Cat. No. 00062107) and Myr-Palm-GFP MDCK cells were maintained in DMEM (Invitrogen, cat. no. 10566016) supplemented with 10% (vol/vol) FBS (Thermo Fisher, Cat. No. 10270106), 1% (vol/vol) Penicillin-Streptomycin (Gibco BRL), and 1% (vol/vol) non-essential amino acids (NEAA) 100X (Invitrogen, ref. 11140050) in cell culture flasks (Falcon) at 37°C and 5% CO2. Encapsulated MDCK monolayers were maintained under the same conditions as above, as well as for live imaging. Cell lines were regularly tested negative for contamination with mycoplasma.

### Microfluidic device fabrication

The microfluidic device (Chip) is composed of two parts. First a 3D printed section, printed with the 3D printer EnvisionTEC Micro Hi-Res Plus, using the resin HTM140V2, with expected Z resolution of 25μm. The following printing parameters (set automatically based on the resin) were used: burn-in range thickness 400 μm, base plate of 300 μm, and exposure time 3000 ms. The printer light intensity is 225 mW/cm. The printed device was washed using ethanol and ddH20 to ensure proper laminar flow and air dried before UV exposition for 30min to ensure full polymerization and increase chip rigidity. The second part, used ensure hydrophobicity and hence proper jet formation while reducing the exit diameter, is a glass capillary (with inner diameter ranging from 50 μm to 200 μm ID) that is added to the tip of the chip. The capillary was pulled and cut to obtain a tip of around 3-4 mm in length and glued on the tip of the microfluidic device with epoxyglue EA M-31CL (Loctite) and left to solidify for 6h at RT. To make the inlets, three 19-gauge stainless steel needles (Terumo, ref.AN-1925R1) were cut into segments 1.5cm long and polished to avoid sharp edges. A small droplet of glue EA M-31CL was spread at the edge of the needles and they were inserted into the inlets of the devices, and the glue was then left to solidify for 24 h at RT. Once the inner glue was solidified, another droplet of glue was added at the inlets edges to ensure proper sealing.

### Microfluidic device operation and cell culture

The working principle of the microfluidic device is explained in detail in (31). In brief, the system is composed of three syringes connected to a pump (Nemesys) for flow rate control, a Matrigel cooling part, the Alginate-charging part system and the microfluidic device. The Chip consists of three coaxial cones inside which three different solutions are injected via the syringes. The outermost cone contains the alginate solution (AL), the intermediate cone contains 300mM sorbitol solution (IS) and the innermost cone contains cells/Matrigel/sorbitol solution (CS) in a ratio of 1:1:2.5 (v/v), with a cell number in the range of 1×10^6 cells. The AL and IS solutions are loaded into two syringes controlled by the pumps for injection into the chip. The CS is injected into a cooling part to maintain Matrigel liquid, and this part is connected to a third syringe containing sorbitol that pushes out the CS into the chip. The volume of Matrigel (Corning, Ref. 354230) used was optimized for a total protein concentration of 0.2mg (optimized to ensure formation of a 3-4μm thick layer of Matrigel). The flow rates are set around 45 mL/h, 40 mL/h and 35 mL/h for AL, IS and CS, respectively, ensuring droplet formation upon exiting the MD, however those rates were varied to alter capsule geometry. Once connected to the pumps the chip is positioned 50-60 cm above a petri dish with a 100mM CaCl2 solution for collection of capsules and 0.1% of TWEEN to reduce surface tension. The alginate charging part and copper ring, both connected to a high voltage (2000V) generator, are used to improve capsule shape and mono-dispersity of size by preventing fusions. The alginate charging part is a glass T connector that has on opposite sides of the T a high voltage wire (coming from the generator) and a tubing containing AL that flows down the connector. The HV wire is coupled to a silver wire (OD 1 mm) that crosses the connector such that it is in contact with the alginate and charges the solution, after which the charged AL then flows into the chip. The copper ring is held below the tip of the MD at a distance of about 1cm and centered with respect to the chip exit flow. The charged formed droplets passing through the copper ring under electrical tension get deflected as they cross the ring, creating a shower-like jet that prevents capsule merging. After 30 min in the calcium bath, capsules are washed and transferred to cell culture medium.

For the cells encapsulated in RGD capsules, the AL was replaced by the 10% or 25% RGD-functionalized alginate solution and the Matrigel was removed from the CS.

For the cells grown on the outer surface of RGD capsules, empty RGD capsules were fabricated by replacing the CS with sorbitol (identical to the IS). After polymerization, the capsules were washed and transferred to a 15mL falcon containing 10 mL of medium. For the seeding, around 1 million suspended cells were added to the falcon. The falcon was placed horizontally in the incubator and gently rotated every 15 minutes for an hour to allow homogeneous cell attachment. The medium was replaced to remove unattached cells, and the cell-loaded capsules were transferred to a petri dish coated with non-adherent solution (to prevent cell and capsule attachment to the bottom) and placed on a slow rocker (to prevent capsule aggregation).

For the hydrogels grown tissues, we used commercially available hydrogels. (Softwell, Easy Coat™ hydrogels bound to 35 mm dishes with a #1.5 glass bottom 0.5 kPa and 4 kPa) The hydrogels where first coated with 1mg/ml bovine plasma fibronectin (Sigma-Aldrich 341631-5mg) in PBS for 30 min at RT prior to cell seeding, at a density of 200 000 cells per well.

### Fluorescent alginate solutions

Labelling of 1% alginate solution with ATTO647N-amine (ATTO-TEC, ref. AD647N-95): 0.25 g Alginate (Protanal LF200FTS, FMS BioPolymer) was dissolved in 25mL 0.1M MES pH 6.0. Next, 5mg ATTO647N-amine dissolved in 200μL DMSO (anhydrous), was added into the tube and mixed, rotating for 30 min. Next, 21.5 mg sulfo-NHS (Sigma, ref. 56485) dissolved in 200μL of 0.1 M MES pH 6.0 was added and let mix for 30min. Finally, 24mg EDC (Sigma, ref.03449) dissolved in 200ul of 0.1M MES pH 6.0 was added and let mix and react overnight at RT. The labeled alginate solution was transferred to a Slide-A-Lyzer™cassette 10K 12-20 mL capacity and let dialyze in milliQ for 2 h, changing milliQ after the 2h and let to dialyze overnight. After dialysis, the labeled alginate was filtered (Acrodisc 25mm Syringe filter with 1um glass finer media, Pall, Life Science). The final concentration of the ATTO647N-labelled alginate was 0.55%, the solution was stored at 4°C. For preparation of 2.5%, 2%, or 1.5% ATTO647N-labeled alginate solutions, 0.27g, 0.165g, 0.22g of alginate powder is mixed with 10 mL milliQ water, respectively. Then, 1mL 0.5% ATTO647N alginate and 10 μl SDS 20% solution (Sigma, ref. 428018) are added, and the mixture is left to rotate overnight at RT. Before use the solutions were spun down at 19,000 rpm for 30 min at 20°C, after which alginate was filtered with a glass-fibre filter before use.

### RGD alginate solutions

Functionalization of alginate with RGD peptide (H-Gly-Gly-Gly-Gly-Arg-Gly-Asp-Ser-Pro-OH PCI-3965-PI) (40): 0.1 g Alginate (Protanal LF200FTS, FMS BioPolymer) was dissolved in 10mL 0.1M MES pH 6.0. Next, 31.8mg sulfo-NHS (Sigma, ref. 56485), 56.3mg EDC (Sigma, ref.03449) and 25mg of RGD were added and let to mix and react overnight at RT. The reaction is then quenched with 18mg hydroxylamine hydrochloride (Sigma, ref. 255580). The RGD-alginate solution was transferred to a Slide-A-Lyzer™cassette 10K 12-20 mL capacity and let dialyze in 4L milliQ with 30 g sodium chloride (Sigma, ref. S9888) for 12 hours, changing milliQ and sodium concentration every 12h (Sodium chloride amounts used: 20g, 10g, 5g). After dialysis, the RGD-alginate was filtered (Acrodisc 25mm Syringe filter with 1um glass finer media, Pall, Life Science), frozen overnight at - 20°C and lyophilized overnight. The dried sponge was stored at -20°C for up to 6 months. For preparation of 2% RGD alginate solutions, 90mg (or 75 mg) of alginate powder is mixed with 5 mL milliQ water. Then, 10mg (or 25 mg) of RGD-alginate sponge and 5 μl SDS 20% solution (Sigma, ref. 428018) are added, and the mixture is left to rotate overnight at RT. The RGD alginate could be used in combination with the fluorescent alginate to generate fluorescent RGD-alginate capsules. Before use the solutions were spun down at 19,000 rpm for 30 min at 20°C, after which alginate was filtered with a glass-fiber filter before use.

### Immunofluorescence

Cell monolayers inside capsules were fixed with 4% (v/v) paraformaldehyde (PFA) (Sigma, ref. F8775) in MEM (not PBS, to avoid dissolving alginate capsule) for 30 min at RT, then permeabilized and dissolved in 100 mM Glycine (Sigma, ref. G8898), 0.1% Triton X-100 (Applichem, ref. A1388) and 1% Gelatin (Sigma, ref. G7765) in 1X PBS for 30 min at RT. Cells were then incubated with prgimary antibody diluted in 1X PBS with 1% Gelatin overnight at 4°C, then washed, and further incubated with secondary antibodies (1:1000) diluted in 1X PBS with 1% Gelatin for 1 h at RT. Primary antibodies used for immunofluorescence staining were: rabbit anti-Vinculin (Thermo Fisher, ref. 700062) and mouse anti-E-Cadherin (BD, ref. 610181). When necessary, monolayers were also counterstained for membrane, f-actin and nuclei using Deep Red Cell Mask (1:1000 Invitrogen C10046), Phalloidin488 (1:40, AlexaFluor488, Thermo Fisher, ref. A12379) and Hoechst 33342 (1:1000, Invitrogen, ref. H3570), respectively. Samples were rinsed 3X with 1X PBS.

### Image acquisition

Capsules were selected between 48 hours and 168 hours post-encapsulation and embedded in 0.4% low-melting agarose (0.04 g in 10 mL) (Sigma, 49414) in a 35 mm glass-bottom dish (Mattek, Part No. P35G-1.0-14-C) to maintain them in the same position for imaging. The agarose was left to solidify for 15 min at RT and 3 mL of RT medium were added for imaging.

For live fluorescence imaging, confocal images of samples were obtained using an inverted Eclipse Ti2 microscope (Nikon) using water immersion objective 40X Nikon CFI APO LWD NIR Objective, 1.15 NA, 0.59 - 0.61 mm WD. During imaging, capsules were maintained at 37 with 5% CO2 for live sample. For each capsule, 3D confocal Z-stacks were acquired with 0.6 μm intervals.

### Image segmentation and processing

Imaris 9.8 software was used to calculate the centroid positions of fluorescent nuclei labelled with Hoechst or H2B-mCherry and calculate their centroid positions. The centroid positions were then used as seeds for the 3D segmentation of the MDCK-II monolayers, labelled with the plasma membrane marker CellMask or Myr-Palm-GFP, using the LimeSeg (41) plugin on Fiji. The centroid positions were also used to reconstruct, via a custom written MATLAB script, the nuclei segmentation performed from each slice using the Stardist plugin for Fiji (42).

MorphoGraphX 1.1 software (43) was used to segment both apical and basal surfaces, to extract cell area and cell-cell junctions’ information. This is done by defining a mesh corresponding to either the apical or the basal surface and projecting the membrane signal contained between 1 um and 3 um away from the mesh. This signal is then segmented via watershed segmentation before the mesh, in addition to the area and junctions’ information, is extracted. The mesh was then used to extract the mean curvature at each points of the mesh using the method from (44).

A custom-written MATLAB script was used to obtain the additional geometrical parameters that where not directly extracted from the software such as the elongation (of apical/basal surfaces and of nuclei) or the pseudo-T1 transition. This was done by linking all the four segmentations together (Apical surface/Basal surface/Cell 3D/Nuclei) and performing data extraction from the meshes, such as the intercellular spacing.

For the immunostaining images quantifications, the freehand election tool in FIJI was used to select the regions of interest (ROI) of the apical and of the lateral F-actin signals in order to quantify the ratio of their intensities, or of the lateral and cytoplasmic E-cadherin signal.

### Data analysis

A custom-written MATLAB script was written to manage and group the 55 000 segmented cells with 24 geometrical parameters. Correlation maps were extracted to identify parameters of interests. For scalar parameters, we used the Pearson correlation coefficient, and for vectorial and nematic parameters we used:

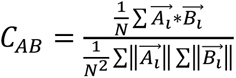 for vectorial parameters

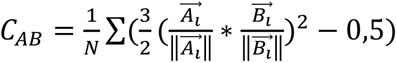 for nematic parameters

All plots were then made using MATLAB inbuilt tools. When indicated on figures, quantities 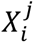 computed for cell *i* in tissue *j*, were normalized by dividing 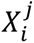 by the average of 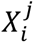 for all cells *i* in tissue *j*.

---

**Supplementary Figure 1:**
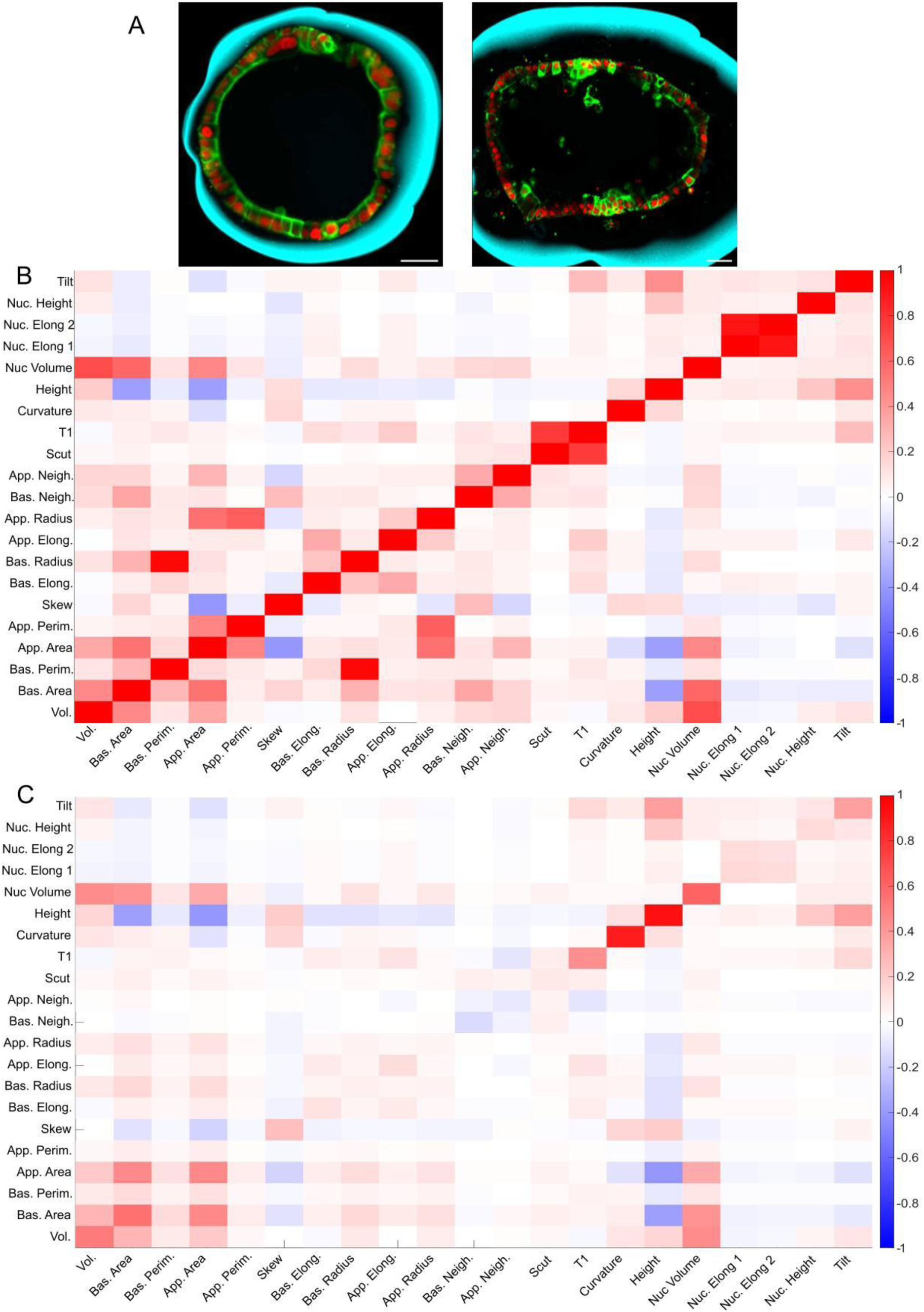
Experimental set up and measured quantities. Tissue/capsule interactions showing confinement and contractile detachment for MP tissues, scale bar 30um (A). Pearson correlation coefficients between pairs of quantities measured on the same cell (B) or on neighboring cells (C) (Matrigel grown tissue, N =47).

**Supplementary Figure 2.**
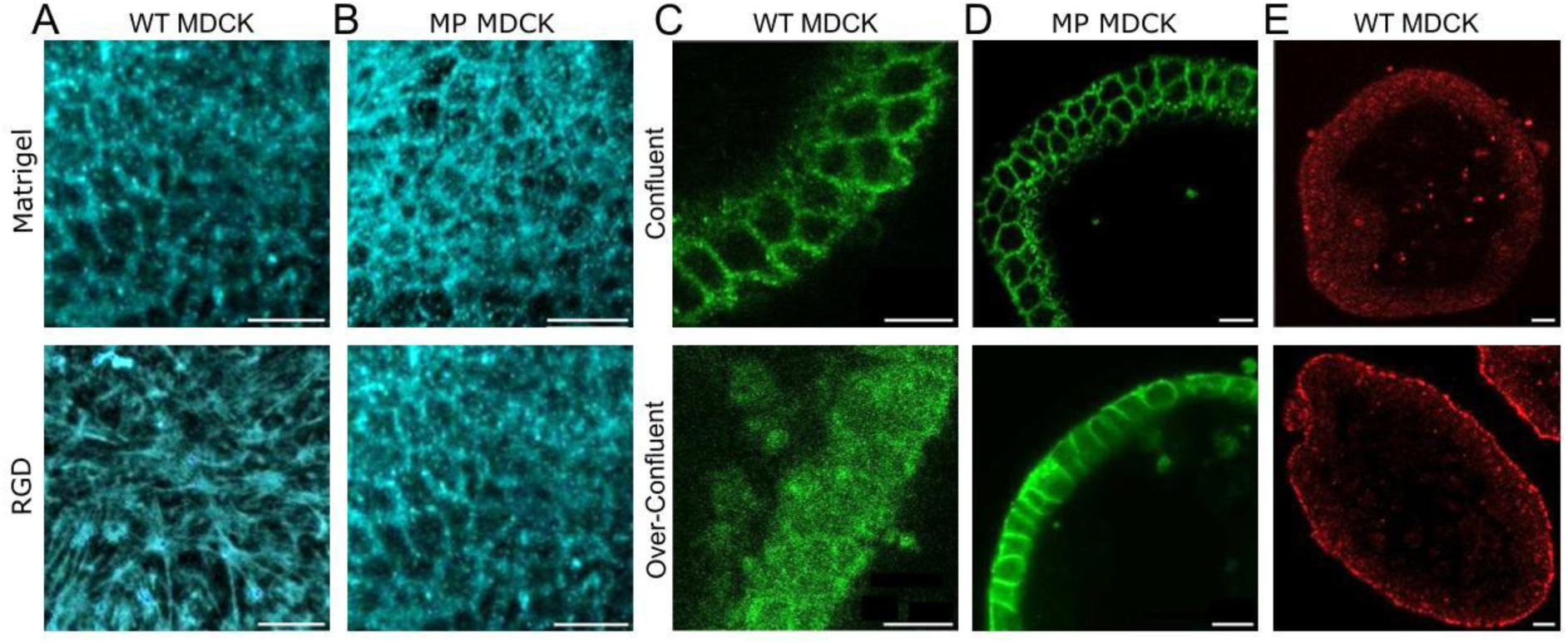
Impact of confinement on tissue organization. Basal actin organization for WT tissues, Matrigel grown (top) and RGD grown (bottom), displaying actin stress fibers on RGD, scale bar 20um (A). Basal actin organization for MP tissues, Matrigel grown (top) and RGD grown (bottom), not displaying actin stress fibers in the RGD grown case, scale bar 20um (B). E-Cadherin in WT MDCK displays a re-localization from cell-cell junctions at normal confluency (top) to the cytoplasm in overconfluent tissues (bottom), scale bar 20um (C). E-Cadherin in MP MDCK displays a re-localization from cell-cell junctions at normal confluency (top) to the cytoplasm in overconfluent tissues (bottom), scale bar 20um (D). Vinculin in WT MDCK displaying an accumulation at the basal side in an overconfluent tissue (bottom) compared to the normal confluency state (top) in a WT Matrigel grown tissue, scale bar 20um(E).

**Supplementary figure 3:**
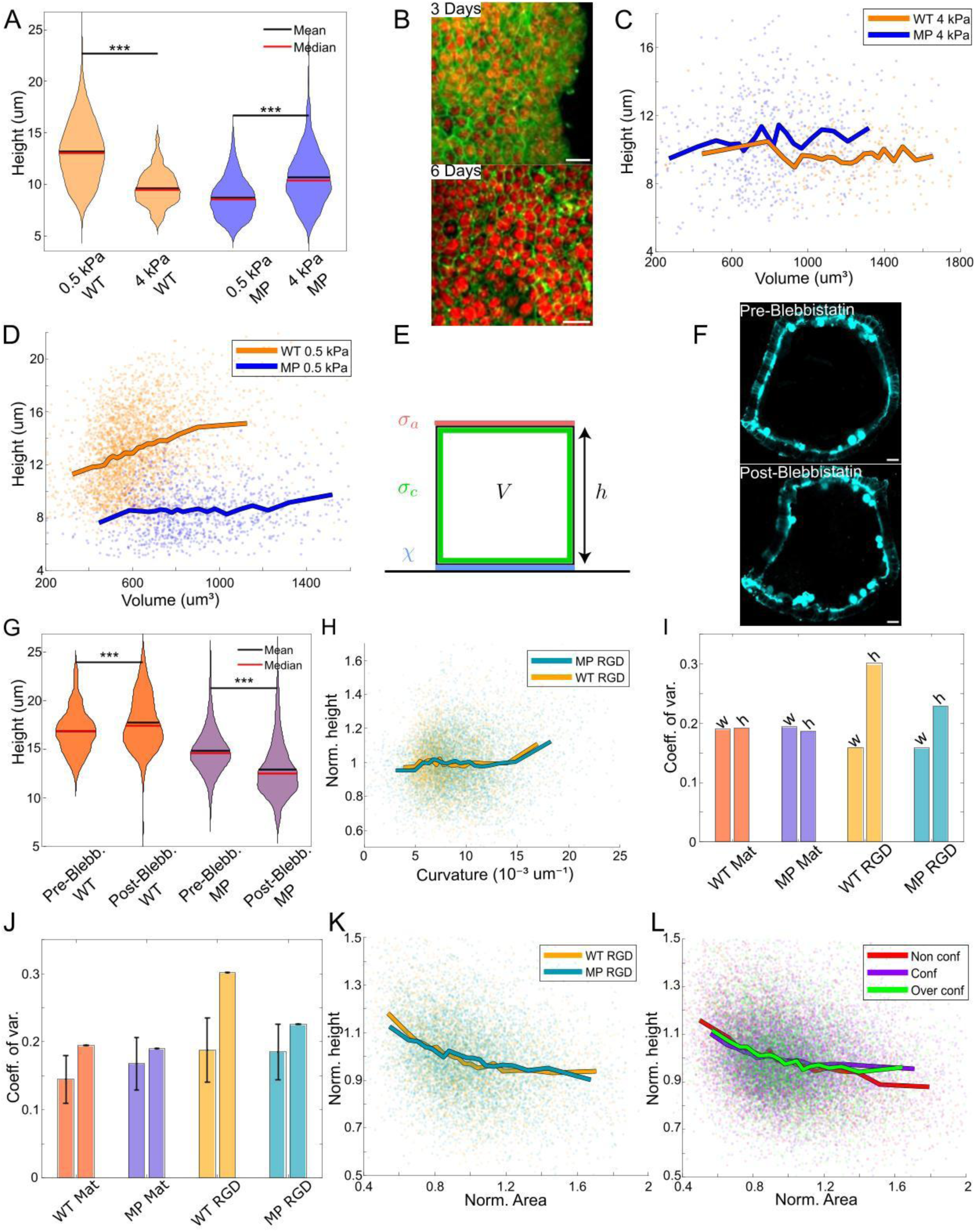
Height control by mechanical parameters Height distribution of hydrogel grown WT and MP tissues, on rigidities of 0.5 kPa and 4kPa. (WT, 0.5 kPa N= 12, 4 kPa N= 10, MP, 0.5 kPa N= 5, 4 kPa N= 5) (A). Hydrogel grown WT tissue, at different confluency, scale bar 10 um, stained for membrane (green) and nuclei (red) (B). Single cell height as a function of volume for WT and MP tissues grown on 4 kPa hydrogels. (WT N=10, MP N=5) (C). Single cell height as a function of volume for WT and MP tissues grown on 0.5 kPa hydrogels. (WT N=12, MP N=5) (D). Illustration of the 3 components considered for the theoretical model, the apical tension *σ*_*a*_, the cortical tension *σ*_*c*_ and the cell-substrate adhesion *χ* (E). Actin staining of capsules prior (Top) and at one hour post blebbistatin treatment (bottom), scale bar 20um (F). Height distribution for WT and MP tissues pre and post blebbistatin treatment (WT N=4, MP N=5) (G). Impact of curvature on tissue height for 25% RGD-alginate capsules (WT: N=13, MP: N=13) (H). Coefficient of variation for cell height and width (*Area*^1/2^), computed across cells of all tissues with identical substrate and cell type (I). Same coefficients of variation for cell height (right bars), compared to the average coefficient of variations of tissues taken individually (left bars, black lines: standard deviation) (J). Normalized height vs normalized area for RGD grown WT and MP tissue (WT: N=13, MP: N=13) (K). Normalized height vs normalized area for Matrigel grown WT in the non-confluent (N=13), confluent (N=11) and overconfluent (N=12) cases (L). Thick lines: average of bins containing equal numbers of elements (C, D, H, K, L). *: p<0.05, **: p< 0.01, ***: p<0.001.

**Supplementary figure 4:**
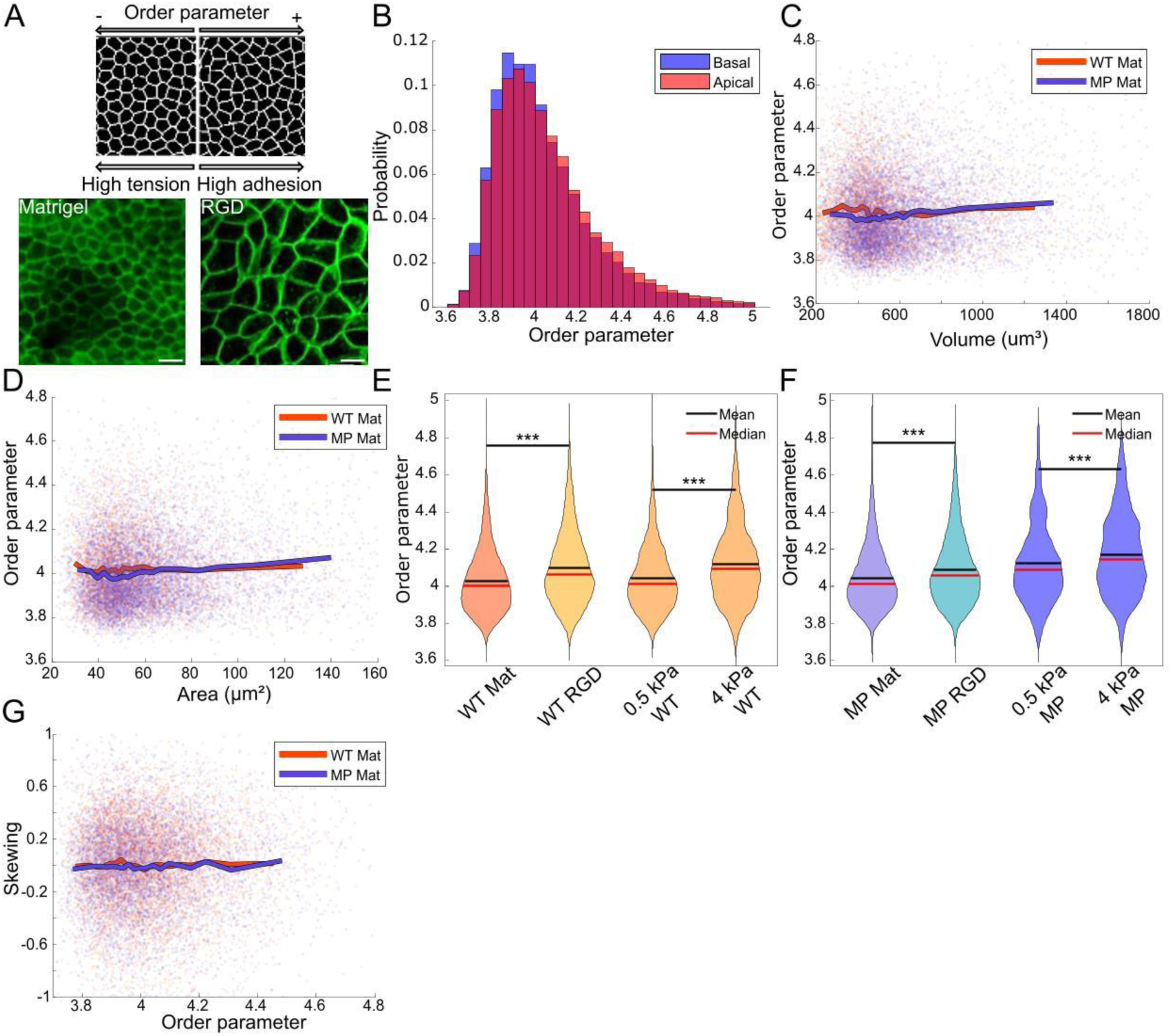
Order parameter quantifications. Illustration of order parameter and corresponding examples (Matrigel grown and RGD grown WT), membrane stained (green), scale bar 10um (A). Order parameter distribution on apical and basal sides, showing minimal differences for Matrigel grown WT (N=20) (B). Order parameter vs cell volume (C) or cell area ((*A*_*a*_ + *A*_*b*_)/2) (D) (WT: N=20, MP: N=27). Order parameter distribution dependence on tissue substrate for WT (Matrigel N=20, RGD N=13, 0.5 kPa Hydrogel N=12, 4kPa Hydrogel N=10) (E) and MP (Matrigel N=27, RGD N=13, 0.5 kPa Hydrogel N=5, 4kPa Hydrogel N=5) tissues (F). Cell skewing vs cell order parameter (WT: N=20 capsules, MP: N=27 capsules) (G). Thick lines: average of bins containing equal numbers of elements (C, D, G). *: p<0.05, **: p< 0.01, ***: p<0.001.

**Supplementary Figure 5:**
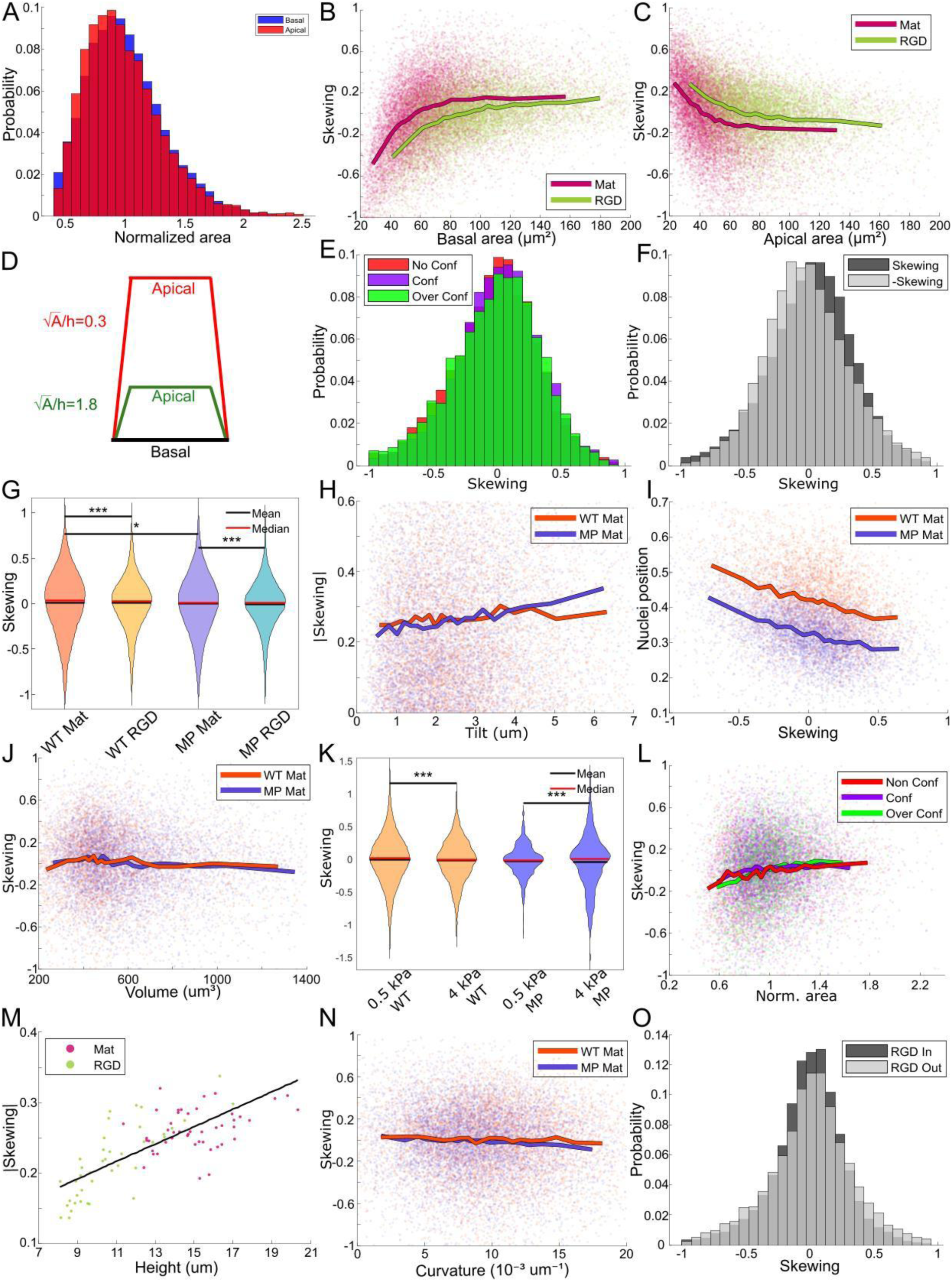
Correlating skewing with other variables Normalized apical and basal area distribution for Matrigel grown WT tissues (N=20) (A). Skewing vs basal area for Matrigel and RGD grown tissues (Matrigel N=47, RGD N=40) (B). Skewing vs apical area for Matrigel and RGD grown tissues (Matrigel N=47, RGD N=40) (C). Illustration of the importance of aspect ratio on the amount of lateral junction tilt for an identical skewing (*S*_*k*_ = 0.5) (D). Distribution of cell skewing depending on tissue state in the non-confluent case (N=13), confluent case (N=11) and overconfluent case (N=12) (E). Skewing (*S*_*k*_) and anti-skewing (−*S*_*k*_) distribution for Matrigel grown tissues (N=47) (F). Skewing distribution based on cell type and substrate (WT Matrigel N=20, RGD N=13, MP Matrigel N=27, RGD N=13)(G). Absolute skewing as a function of cell tilt for Matrigel grown tissues (WT: N=20, MP: N=27) (H). Nuclei position along the apico-basal axis (0: basal side, 1:apical side) vs skewing, for Matrigel grown tissues (WT: N=20, MP: N=27) (I). Cell skewing as a function of cell volume (WT: N=20, MP: N=27) (J). Skewing distribution for Hydrogel grown tissues of different rigidities (WT 0.5 kPa Hydrogel N=12, 4kPa Hydrogel N=10, MP 0.5 kPa Hydrogel N=5, 4kPa Hydrogel N=5) (K). Skewing distribution as a function of normalized area for WT Matrigel grown tissues at different tissue states, in the non-confluent (N=13), confluent (N=11) and overconfluent (N=12) cases (L). Capsule average absolute skewing as a function of capsule average cell height (Matrigel N=47, RGD N=26, black line: linear fit) (M). Skewing vs curvature for Matrigel grown WT and MP (WT: N=20, MP: N=27) (N). Skewing distribution for concave (In) and convex (Out) RGD-grown tissues (RGD 25% In N=25, RGD Out N=13) (O). Thick lines: average of bins containing equal numbers of elements (B, C, H, I, J, N). *: p<0.05, **: p< 0.01, ***: p<0.001.

**Supplementary Figure 6:**
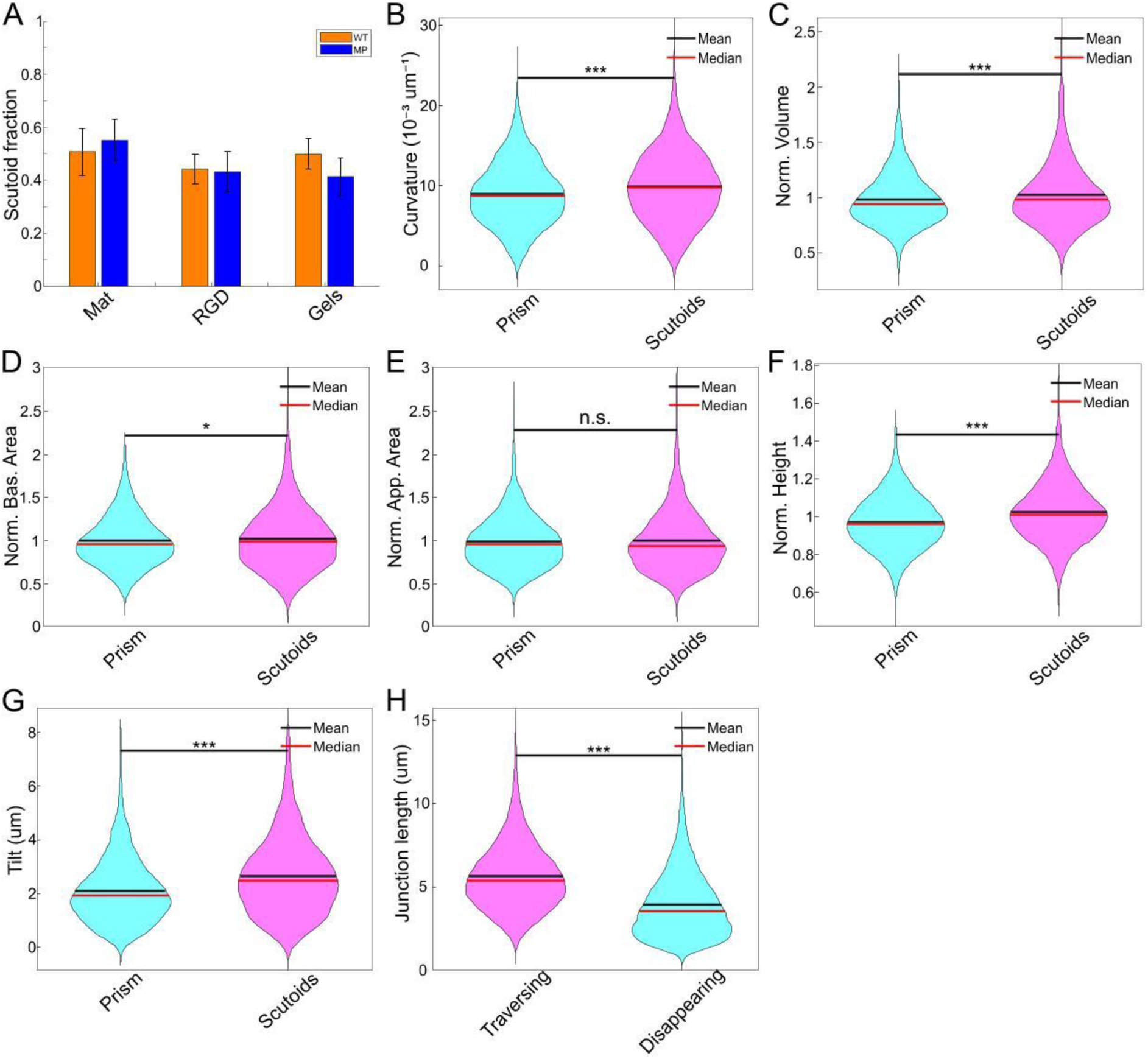
Differences in organization between prisms and scutoids. Fraction of scutoids present in the different tissues (WT Matrigel N=20, RGD N=13, Hydrogel N=22, MP Matrigel N=27, RGD N=13, Hydrogel N=9) (A). Distribution, for prisms and scutoids separately, of substrate curvature (B), normalized volume (C), normalized basal area (D), normalized apical area (E), normalized height (F), and tilt (G). Distribution (for all cells taken together) of length of junctions that either traverse the entire cell along the apical-basal axis (Traversing), or disappears along the apical-basal axis (Disappearing) (H). Matrigel grown tissues (N=47) (B-H). *: p<0.05, **: p< 0.01, ***: p<0.001.

## Notes

### Competing Interest Statement

The authors have declared no competing interest.

